# Effects of decreased Rac activity and malignant state on oral squamous cell carcinoma

**DOI:** 10.1101/540591

**Authors:** Hani Al-Shareef, Yudai Matsuoka, Mikihiko Kogo, Hirokazu Nakahara

**Affiliations:** Department of Oral & Maxillofacial Surgery, Osaka City University Graduate School of Medicine, Osaka, Osaka, Japan; The First Department of Oral & Maxillofacial Surgery, Osaka University Graduate School of Dentistry, Suita, Osaka, Japan

**Keywords:** Rac, apoptosis, JNK, anti-cancer therapy

## Abstract

Rac proteins, members of the Rho family of small GTP-binding proteins, have been implicated in transducing a number of signals for various biological mechanisms, including cell cytoskeleton organization, transcription, proliferation, migration, and cancer cell motility. Among human cancers, Rac proteins are highly activated by either overexpression of the genes, up-regulation of the protein, or by mutations that allow the protein to elude normal regulatory signaling pathways. Rac proteins are involved in controlling cell survival and apoptosis. The effects of Rac inhibition by the Rac-specific small molecule inhibitor NSC23766 or by transfection of dominant negative Rac (Rac-DN) were examined on three human-derived oral squamous cell carcinoma cell lines that exhibit different malignancy grades, OSC-20 (grade 3), OSC-19 (grade 4C), and HOC313 (grade 4D). Upon suppression of Rac, OSC-19 and HOC313 cells showed significant decreases in Rac activity and resulted in condensation of the nuclei and up- regulation of c-Jun N-terminal kinase (JNK), leading to caspase-dependent apoptosis. In contrast, OSC-20 cells showed only a slight decrease in Rac activity, which resulted in slight activation of JNK and no change in the nuclei. Fibroblasts treated with NSC23766 also showed only a slight decrease in Rac activity with no change in the nuclei or JNK activity. Our results indicated that apoptosis elicited by the inhibition of Rac depended on the extent of decreased Rac activity and the malignant state of the squamous cell carcinoma. In addition, activation of JNK strongly correlated with apoptosis. Rac inhibition may represent a novel therapeutic approach for cancer treatment.

## Introduction

Rac proteins have been implicated in transducing a variety of signal pathways that are essential for cell function [1–4]. Upon stimulation by growth factors, Rac proteins are activated through a tightly regulated guanosine diphosphate/guanosine triphosphate (GDP/GTP) cycle. Activated Rac proteins are a key regulator of a number of cell activities, such as cell organization of the cytoskeleton, transcription, cell proliferation, cell migration, and cancer cell motility [5,6]. Rac proteins are also involved in controlling cell survival and apoptosis [7]. Activation of the Rac family induces a strong signal to activate cancer cells and related fibroblasts in the apoptosis mechanism [8–11]. In most human cancers, Rac proteins are highly activated by either overexpression of the gene, up-regulation of the protein, or mutations that allow the protein to elude normal regulatory signaling pathways [12–15]. However, the mechanisms of Rac GTPases and their relation to apoptosis require additional studies, which could contribute toward their development as therapeutic agents in cancer treatment.

Activation of c-Jun N-terminal kinase (JNK) by various stimulatory signals results in an apoptotic response via a number of its substrate effectors. Many studies have indicated that Rac is an upstream activator of JNK/c-Jun signaling [9,16,17] and others have recently reported that Rac can also suppress the JNK signaling pathway [10–18].

In the current study, we investigated the role of Rac in the malignant oral squamous carcinoma cell lines OSC-20, OSC-19, and HOC313. We found that apoptosis induced by the inhibition of Rac was dependent on the extent of decreased Rac activity and the malignant state of the squamous cell carcinoma. We also determined that activation of JNK strongly correlated with the induced apoptosis.

## Results

### Characterization of three human-derived oral squamous cell carcinoma cell lines, OSC-20, OSC-19, and HOC313 cells

The three cell lines used in the current study, OSC-20, OSC-19, and HOC313 cells, were derived from human oral squamous cell carcinomas and each demonstrates different malignancy according to Yamamoto-Kohama’s (Y-K) classification [19]. OSC-20 cells are classified as grade 3, OSC-19 as grade 4C, and HOC313 as grade 4D. To characterize each cell type, the cells were cultured on microscope coverslips and stained with rhodamine-phalloidin. As shown in Fig 1A, OSC-20 cells formed lamellipodia, OSC-19 cells formed filopodia, and HOC313 cells formed abundant horizontal growth stress fibers with microspikes. To further investigate the characteristics of each cell, we then examined the cytoskeletal proteins of three cell lines. OSC-19 cells stained positive for E-cadherin and cytokeratin (Fig 1B). On the other hand, HOC313 stained positive for N-cadherin and vimentin (Fig 1B). As shown in Fig 1B, OSC-20 cells showed staining for E-cadherin and N-cadherin, cytokeratin, and vimentin. These results indicated that OSC-19 cells had epidermal characteristics and HOC313 cells had mesenchymal characteristics. OSC-20 cells were characterized as an intermediate between OSC-19 and HOC313 cells.

**Fig 1.**
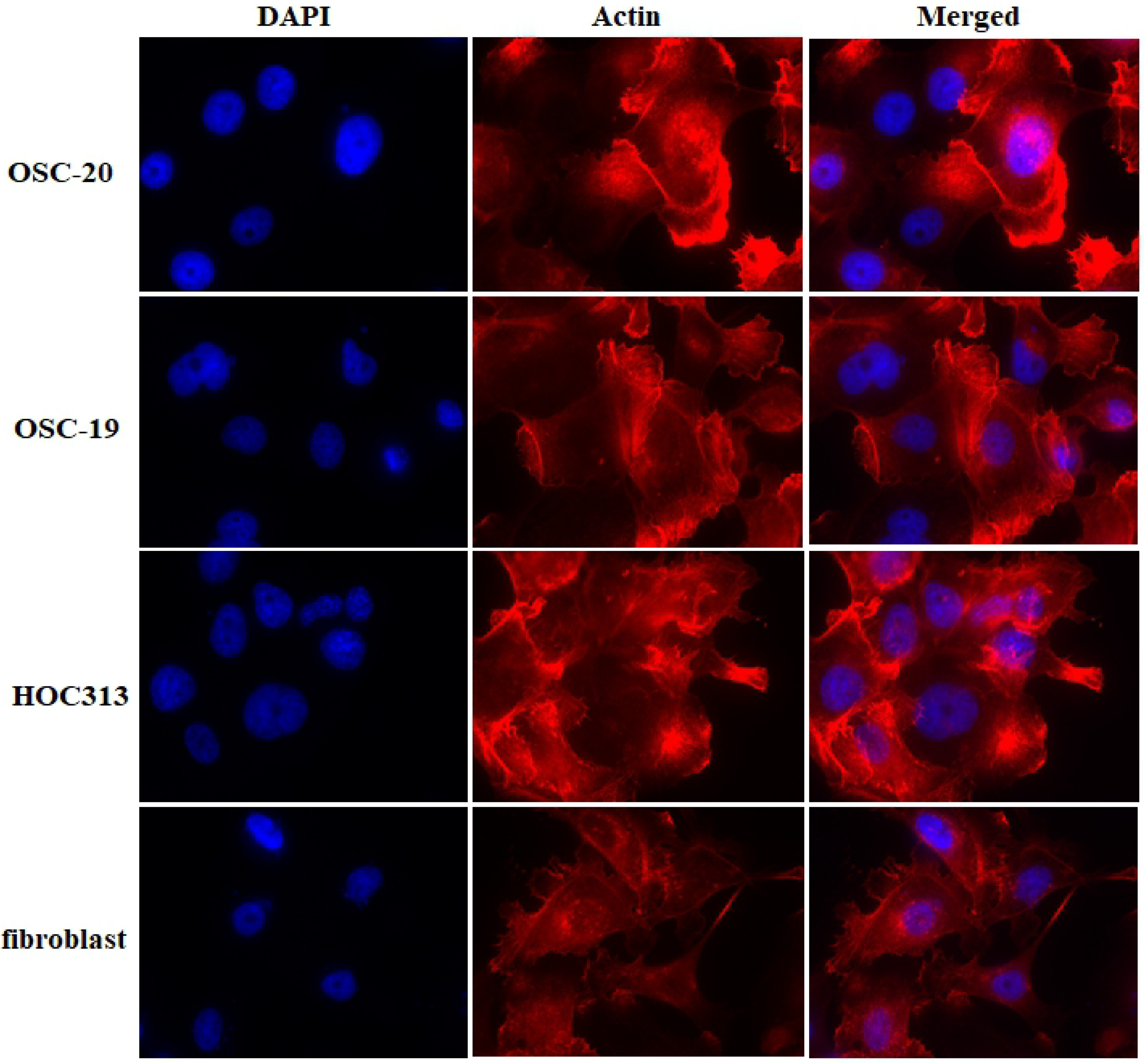
Characteristics of three human-derived oral squamous cell carcinoma cell lines, OSC-20, OSC-19, HOC313 cells, and fibroblasts. All cells were seeded onto coverslips and grown in DMEM containing 10% fetal bovine serum and 5% Nu-serum Growth Medium Supplement for 24 h and then stained with rhodamine-phalloidin to compare morphology.

### Inhibition of Rac activity induced cell death in OSC-19 and HOC313 cells but not in OSC-20 cells or fibroblasts

Based on the molecular evidence that Rac activity increases as a mechanism of cancer, we hypothesized that inhibition of Rac might be detrimental to tumor cells. To test this hypothesis, we evaluated the effects of the selective Rac1 inhibitor NSC23766 on OSC- 20, OSC-19, and HOC313 cells. Impressively, OSC-19 and HOC313 cell lines treated with 100 μM NSC23766 for 24 h showed morphological changes indicative of cell death whereas the OSC-20 cell line failed to show morphological changes associated with cell death (Fig 2). Furthermore, similar to the OSC-20 cells, fibroblasts treated with 100 μM NSC23766 for 24 h also showed no cell death related morphology (Fig 2). We performed western blot analysis of pull-down assays for the whole Rac protein and a Rac effector domain to evaluate any changes or association differences in Rac activity among the three SCC cell lines and fibroblasts. Based on the results, the Rac expression in general was obvious in the OSC-20, OSC-19, and HOC313 cell lines and the Rac pull-down assay revealed a high level of Rac activity, more than that in the fibroblasts under the untreated condition (Fig 3). These data suggested that Rac was overexpressed and had a higher level of activity in OSC-20, OSC-19, and HOC313 cell lines compared with those in the fibroblasts. Moreover, a significant decrease in Rac activity was also observed in all the cell lines after treatment with NSC23766 (Fig 3). However, the extent of decreased Rac activity in OSC-19 and HOC313 cells was greater than that of OSC-20 cells and fibroblasts (Fig 3).

**Fig 2.**
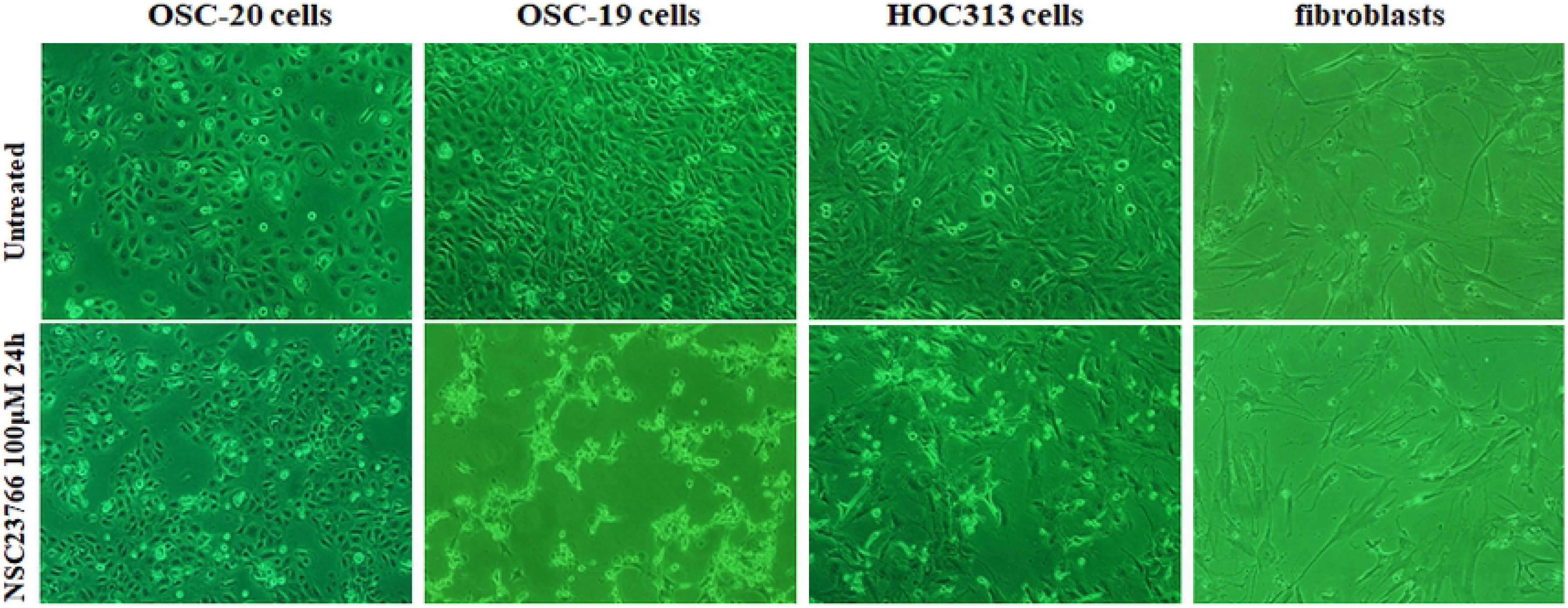
Phase-contrast micrographs demonstrating the effects of the selective Rac1 inhibitor NSC23766 on OSC-20, OSC-19, HOC313, and fibroblast cells. All cells were treated with 100 μM NSC23766 for 24 h. OSC-19 and HOC313 cells showed morphological changes associated with cell death, in contrast to OSC-20 cells and fibroblasts, which did not demonstrate morphological changes.

**Fig 3.**
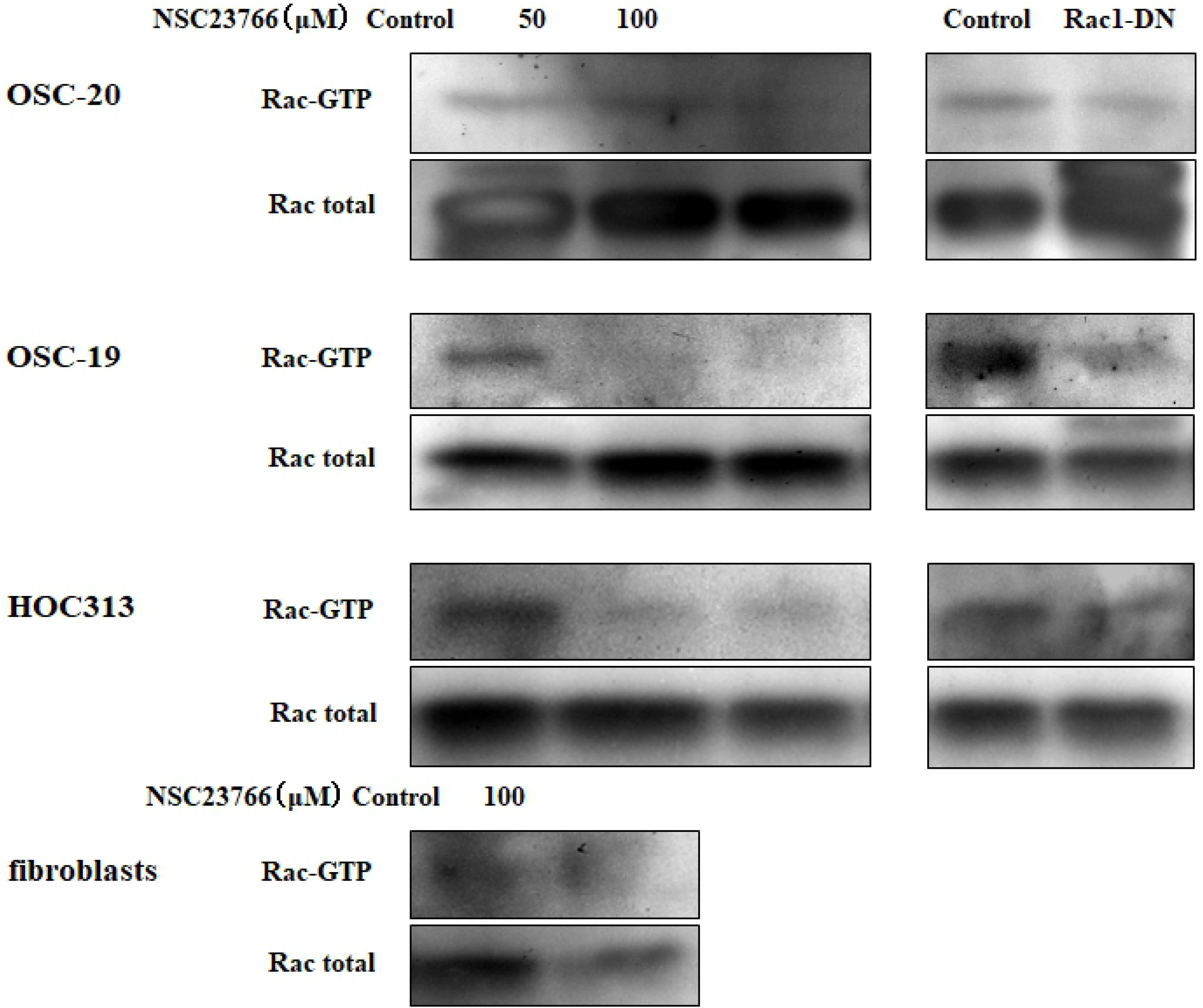
Western blot analyses of pull-down assays for the whole Rac protein and a Rac effector domain. The extent of decreased Rac activity in OSC-19 and HOC313 cells was more significant than that in OSC-20 cells and fibroblasts. The three human-derived oral squamous cell carcinoma cell lines (OSC-20, OSC-19, HOC313), and fibroblasts were treated with 0, 50, or 100 μM selective Rac1 inhibitor NSC23766 and incubated them for 9 h in serum followed by the transient transfection with the dominant negative Rac (Rac-DN) for 48 h. Pull down the GTP-loaded Rac from the total protein lysates was performed using a Rac1 Activation Kit (GST-human Pak1-PBD) according to the manufacturer’s instructions.

### The type of cell death elicited by inhibition of Rac activity was apoptosis

After treating the SCC cells and fibroblasts with 100 μM NSC23766 to inhibit Rac activity and staining the nuclei with 4′,6-diamidino-2-phenylindole (DAPI) stain, we examined the cells under a microscope and checked their structure to determine whether the cell death had occurred due to necrosis or apoptosis. We observed morphological characteristics of apoptosis, such as cell shrinkage, nuclear condensation, and fragmentation in the OSC-19 and HOC313 cells (Fig 4A). In contrast, these findings were not seen in the OSC-20 cells or fibroblasts after they were treated with the same procedure (Fig 4A). We also performed a cell death detection enzyme-linked immunosorbent assay (ELISA) to confirm our results regarding the apoptotic cell death (Fig 4B). The results revealed a significant 2–6-fold increase in apoptosis in OSC-19 and HOC313 cells after treatment with NSC23766 compared to the control cells, OSC-20 cells, and fibroblasts after treatment (p < 0.05). These results suggested that the inhibition of Rac activity by the Rac-specific small molecule inhibitor NSC23766 had a significant role in inducing apoptosis in the OSC-19 and HOC313 cells. To extend these findings, we performed additional investigations into the mechanism of inhibition of Rac activity. Briefly, the cells were transfected with an expression vector encoding a Myc-tagged Rac dominant negative mutant (Rac-DN) and the expression levels were measured in the treated cells. The Rac-DN transformed cell showed cell shrinkage, nuclear condensation, and fragmentation in the cell nuclei (Fig 4A). To confirm our results regarding apoptotic cell death in the transformed cells, we then performed cell death detection ELISA assay (Fig 4B). The amount of cell death resulting from apoptosis for the cells treated with Rac-DN appeared less than that observed for the cells treated with 100 μM NSC23766. Analytical evaluation of the results revealed a negative correlation, which was reasonable since the transfection efficiency was approximately 30–40% and a small amount of Rac activity was detected in Rac pull-down assays of cells that underwent the transient expression with Rac-DN (Fig 3). Overall, these results suggested that the inhibition of Rac activity was likely to lead to the cell apoptosis in OSC-19 and HOC313 cells, which was in contrast to OSC-20 cells or fibroblasts in which no cell apoptosis was observed.

**Fig 4.**
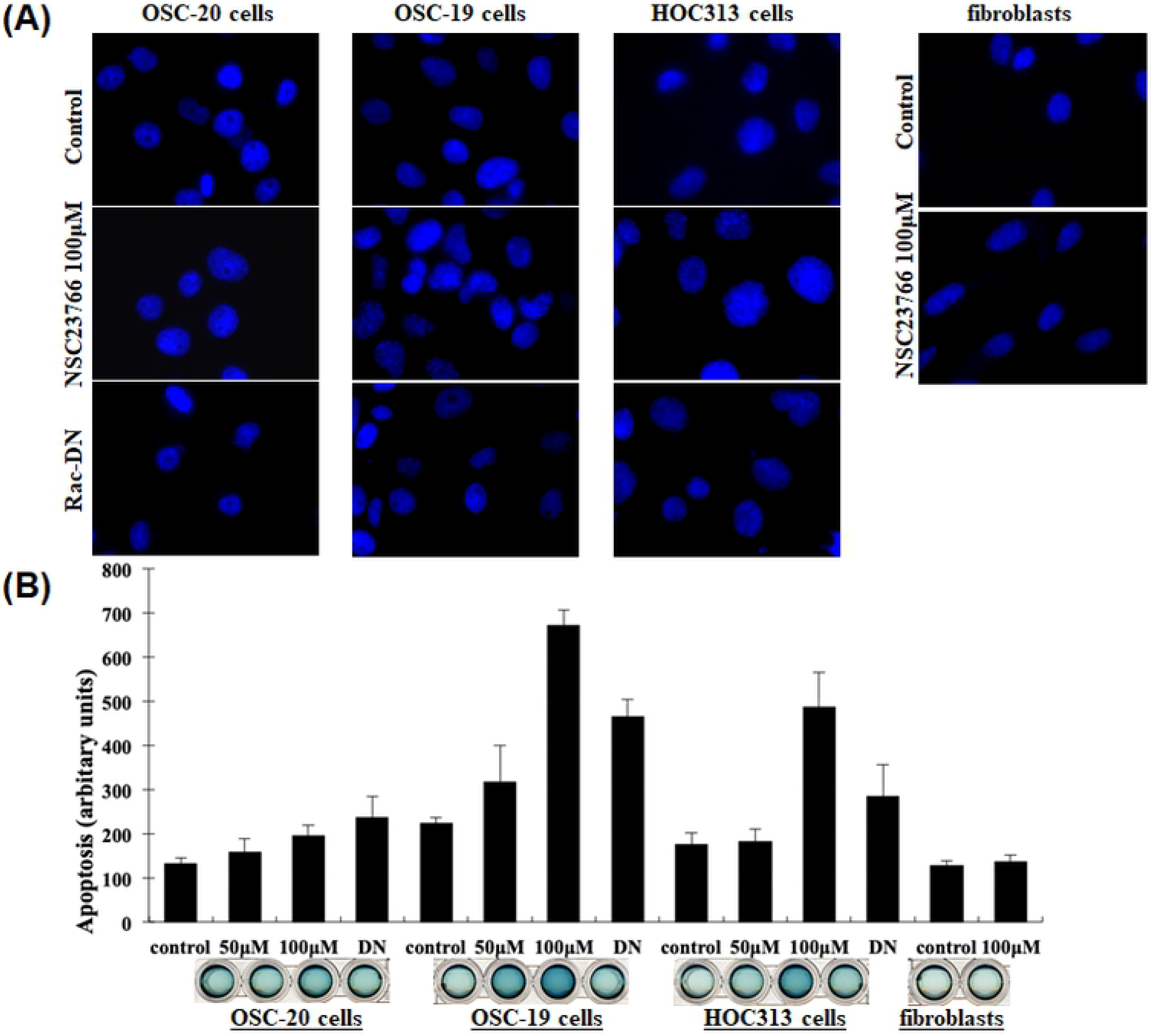
DAPI staining of nuclei and cell death detection ELISA assay. (**A**) The three human-derived oral squamous cell carcinoma cell lines, OSC-20, OSC-19, and HOC313, and human fibroblasts were seeded onto round glass coverslips glass (105 cells). The cells were experimentally treated and then fixed in 4% paraformaldehyde, washed, and stained with DAPI for 1 h at room temperature. The cells were treated with selective Rac1 inhibitor NSC23766 (100 μM) for 9 h, then underwent transient transfection with an expression vector of a dominant negative Rac (Rac-DN). Fibroblasts were treated with NSC23766 (100 μM) under the same conditions. (**B**) A cell death detection ELISA assay was performed according to the manufacturer’s instructions to confirm the results regarding the apoptotic cell death.

### Hyper-phosphorylation of JNK correlated with apoptosis induced by the inhibition of Rac

Studies have suggested that Rac acts as an upstream activator of JNK/c-Jun signaling [9,16,17], while others have suggested that Rac can also suppress the JNK pathway [10,18]. To expand our current studies related to the inhibition of Rac activity and its relation to apoptosis in OSC-19 and HOC313 cells, we investigated downstream signaling of the Rac pathway. We briefly inhibited the Rac activity by treating the cells with 100 μM NSC23766 for 9 h and the examined the phosphorylation of JNK. We detected hyper-phosphorylation of JNK, as assessed by a phospho-specific antibody (Fig 5A). In contrast, OSC-20 cells treated with 100 μM NSC23766 under the same conditions showed only slight activation of JNK. Furthermore, there was no difference detected in JNK activity for fibroblasts after being treated under the same circumstances (Fig 5A).

**Fig 5.**
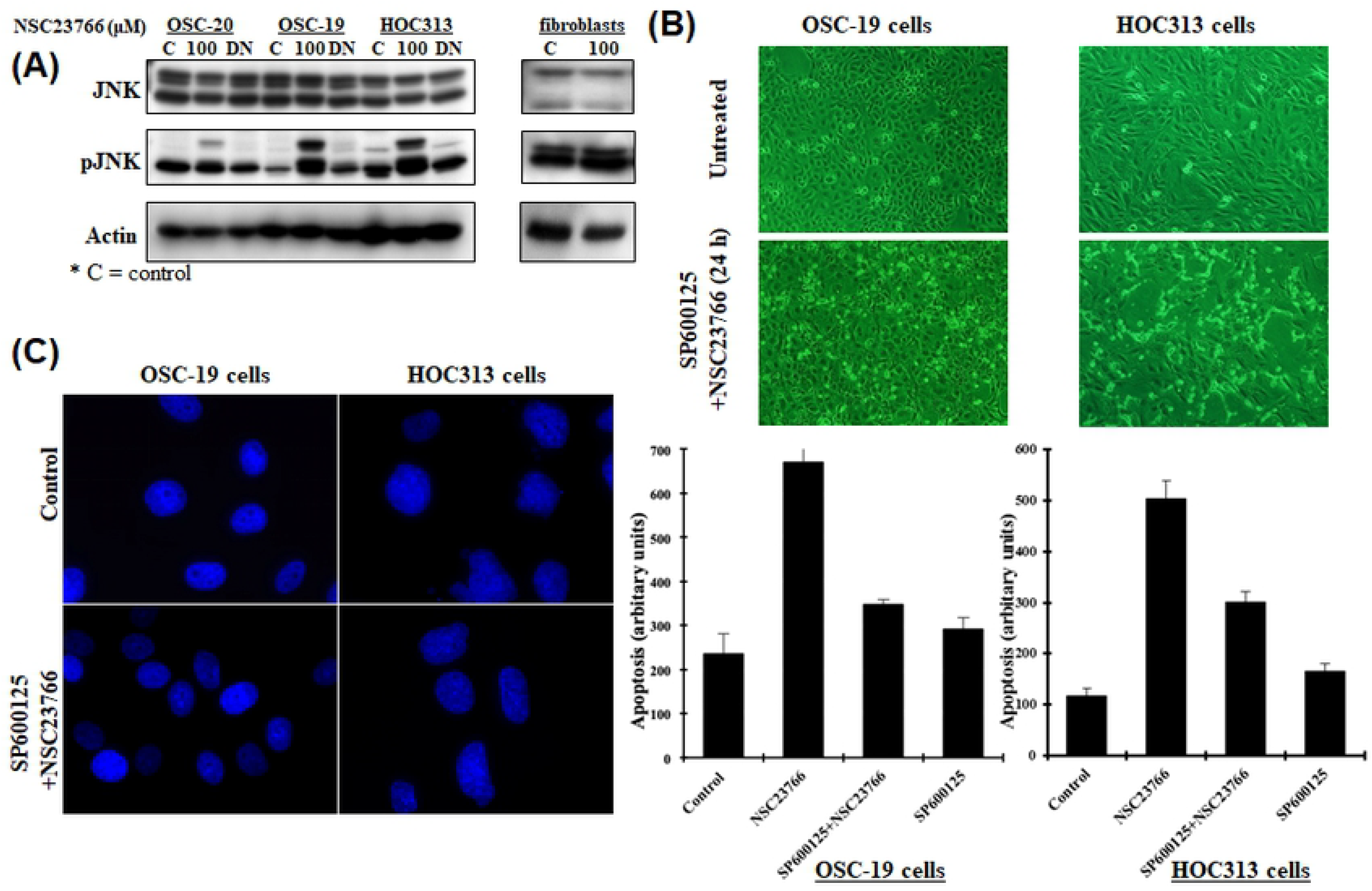
(**A**) Western blot analysis of JNK using anti-JNK and pJNK polyclonal antibodies and a β-actin monoclonal antibody. Rac activity was inhibited by treating the human-derived oral squamous cell carcinoma cell lines OSC-19, OSC-20, and HOC313 with 100 μM selective Rac1 inhibitor NSC23766 for 9 h followed by the transient transfection of the three cell lines with a dominant negative Rac (Rac-DN) for 48 h. The phosphorylation of JNK was examined. Fibroblasts were treated with NSC23766 (100 μM) under the same conditions. (**B**) Phase-contrast micrographs. OSC-19 and HOC313 cells were pretreated with JNK-specific inhibitor (20 μM SP600125) for 1 h before treatment with 100 μM NSC23766 for 24 h. (**C**) DAPI staining of nuclei to detect morphological changes. OSC-19 and HOC313 cells were seeded onto coverslips, the cells were experimentally treated, and fixed in 4% paraformaldehyde, washed and stained with DAPI for 1 h at room temperature. The DAPI-stained cells were then pretreated with JNK-specific inhibitor (20 μM SP600125) for 1 h before treatment with the 100 μM NSC23766 for 9 h. Cell death was detected by an ELISA assay according to the manufacturer’s instructions to confirm our results regarding the apoptotic cell death.

To further confirm whether the inhibition of Rac activity in relation to apoptosis was mediated through a JNK-upregulation mechanism, we pretreated the OSC-19 and HOC313 cells with the JNK-specific inhibitor SP600125 (20 μM) for 1 h prior to treating the cells with NSC23766 [19-20] and evaluated the effect on apoptosis. The cells failed to show hallmarks of cell-death, even after 24 h of incubation with 100 μM NSC23766 (Fig 5B). Treating the cells with SP600125 prevented the condensation and fragmentation of the cell nuclei and analysis using the cell death detection ELISA assay showed that apoptosis was decreased by approximately 50% (Fig 5C). In conclusion, these results suggested that the activation of JNK was more effective after inhibition of Rac activity in OSC-19 and HOC313 cells compared to OSC-20 cells and fibroblasts and was necessary for the induction of apoptosis.

## Discussion

In particular, several lines of evidence indicate that Rac proteins play crucial roles in several aspects of cell survival and apoptosis [7,21]. Therefore, inhibition of Rac using specific inhibitors may allow for the modulation of apoptosis [6]. In the current study, a couple key included: (a) inhibition of Rac activity and its mediated downstream signaling likely led to the cell apoptosis observed for the OSC-19 and HOC313 cells, but not for the OSC-20 cells or fibroblasts; (b) the activation of JNK signaling in OSC-19 and HOC313 cells was associated with the inhibition of Rac activity and thereby to apoptosis. Consequently, activation of the Rac pathway may indicate the preservation of cancer cell progression and the selective inhibition of Rac activity may trigger apoptosis in cancer cells. Therefore, inducing apoptosis in cancer cells by inhibiting Rac activity may be exploited as a novel treatment for cancer. However, Rac activity was more readily suppressed in OSC-19 and HOC313 cells compared to OSC-20 cells and fibroblasts suggesting potential variation in response to treatment.

One mechanism that requires additional elucidation is the regulation of JNK activity after suppression of Rac. Many studies have indicated that Rac acts as an upstream activator of JNK signaling in certain cells [9,16,17], while others have reported that Rac can suppress the JNK pathway [10,18]. In general, whether Rac positively or negatively regulates the JNK pathway may primarily depends on the specific cancer cell type.

It is important to emphasize that the hyper-phosphorylation of JNK was activated by the inhibition of Rac activity in OSC-19 and HOC313 cells by treating the cells with 100 μM NSC23766, but this activation of JNK did not occur in OSC-20 cells or fibroblasts. Moreover, the suppression of JNK activity by treating the OSC-19 and HOC313 cells with a JNK-specific inhibitor induced similar effects on apoptosis as the direct inhibition of Rac. Taken together, these findings suggest that the activation of JNK was more effective after treatment to inhibit Rac activity in OSC-19 and HOC313 cells compared to OSC-20 cells and fibroblasts and that JNK activation was necessary for induction of apoptosis.

In conclusion, results from our study suggest that the inhibition of Rac activity resulted in the hyper activation of JNK, which led to apoptosis in OSC-19 and HOC313 cells, but not in OSC-20 cells or fibroblasts. Additional studies are needed to develop a better understanding of the apoptosis mechanism relative to the suppression of Rac activity and its related regulatory signal pathways. The end result of these efforts may lead to the development and design new strategic therapeutic approaches in the treatment of oral squamous cell carcinoma.

## Materials and Methods

### Antibodies and chemicals

Anti-Rac1 (23A8) and anti-β-actin (MAB 1501) monoclonal antibodies were purchased from [Upstate Biotechnology and Chemicon International, respectively (now Merck KGaA, and its affiliates), Darmstadt, Germany]. Both the anti-JNK and anti-phospho-JNK (Thr183/Tyr185) polyclonal antibodies were purchased from R&D Systems, Inc. (Minneapolis, MN, USA). E-cadherin (36/E-Cadherin) and N-cadherin (32/N-Cadherin) were obtained from BD Biosciences (Becton Dickinson, Franklin Lakes, New Jersey, USA). Cytokeratin (AE1/3) and vimentin (V9) were purchased from Dako Agilent Technologies, Inc. (Santa Clara, CA, USA). Secondary antibodies were purchased from Cell Signaling Technology, Inc. (Danvers, MA, USA). The complementary DNA (cDNA) of the dominant negative N17Rac1 (Rac-DN), which encoded for a mutated amino acid 17 from Thr to Asn was provided by Dr. A. Hall (University College London, Laboratory for Molecular Cell Biology, UK). The expression plasmid was constructed as previously described [22].

### Cell culture and cell treatment

Three human-derived oral SCC cell lines, OSC-20, OSC-19, and HOC313 were previously established and used in the study. OSC-20 cells are classified as grade 3 using the Y-K classification regarding mode of invasion, OSC-19 cells are classified grade 4C, and HOC313 cells are classified as grade 4D. Fibroblasts cells were isolated from the lip skin of adult patients. All clinical studies were approved by the Ethics Committees of Osaka City University Hospital and Osaka University Dental Hospital. The cancer cells including the fibroblasts were cultured in Dulbecco’s modified Eagle’s medium (DMEM) supplemented with 10% fetal bovine serum, 5% Nu-serum Growth Medium Supplement (Oscient Pharmaceuticals Corp., Waltham, Massachusetts, USA), and 2 mM L-glutamine at 37°C and 5% CO2 humidified atmosphere. The cells were typically passaged when they reached 85–90% confluency. The cells were seeded into 6-well cell culture plates with 2 ml growth medium and any treatment was applied the following day. All inhibitors were added 1 h prior to treating the cells with NSC23766. Once the NSC23766 was added, the cells were incubated for 9 h. The dimethyl sulfoxide (DMSO) concentrations were maintained at equal concentrations in the control cells and in those receiving the inhibitors in the media. The DMSO concentration never exceeded 0.1%. The three SCC cell lines were transiently transfected with the expression plasmids using FuGENE HD transfection reagent (Roche Molecular Systems, Inc., Upper Bavaria, Germany) according to the manufacturer’s instructions as described in the online link.

### Rac pull-down assay and western blotting

The three SCC cell lines (OSC-20, OSC-19, and HOC313) and the fibroblasts were treated with 0, 50, or 100 μM of NSC23766 and incubated for 9 h in serum. After treatment, the cells were transiently transfected with Rac-DN for 48 h. The cells were then lysed with 0.3 ml of a lysis buffer (25 mM Tris, pH7.5; 150 mM NaCl; 5 mM MgCl2; 1% NP-40; 1 mM DTT; and 5% glycerol), mixed well, and incubated at 4°C for 5 min. The lysates were carefully clarified, the protein concentrations normalized, and the GTP-loaded Rac pulled down from the total protein lysates using a Rac1 Activation Kit (GST-human Pak1-PBD, Thermo Fisher Scientific, Waltham, MA, USA) according to the manufacturer’s instructions. The precipitates were washed and boiled and the proteins separated by 10% sodium dodecyl sulfate polyacrylamide gel electrophoresis (SDS-PAGE). The proteins were transferred from the gels onto nitrocellulose membranes and quantified by anti-Rac1 immunoblotting.

For western blot analysis, a RIPA Lysis Buffer system (Santa Cruz Biotechnology, Inc., Dallas, Texas, USA) was used to lyse the cells and generate the cell extracts. The lysates were clarified, the protein concentrations adjusted to standards amounts, and boiled in 3× sample buffer. The samples were separated by SDS-PAGE, the proteins were transferred onto nitrocellulose membranes. The membranes were subsequently blocked with 5% membrane blocking agent (skim milk) for 1 h incubated with the appropriate antibodies and then visualized using Amersham ECL Prime Western Blotting Detection Reagent (GE Healthcare, Chicago, IL, USA).

### Immunofluorescence staining

To detect the morphological changes of the nuclei, the cells were stained with DAPI. Briefly, OSC-20, OSC-19, HOC313, and fibroblast cells were seeded (105 cells) onto round glass coverslips, #1.5 thickness, 18 mm in 12-well tissue culture plates. The cells were experimentally treated as described above and then fixed in 4% paraformaldehyde, washed, and incubated with DAPI stain for 1 h at room temperature. After DAPI staining, the cells were stained with rhodamine-phalloidin for 1 h at room temperature, washed, and the cells adhering to the coverslips were mounted onto glass microscope slides using mounting medium. An Axiovert 200M Inverted microscope (Carl Zeiss, Germany) was used to visualize the cells and to capture the resulting fluorescent images.

### Detection of apoptosis

To detect apoptosis, we used a cell death detection ELISA (Roche Molecular Systems, Inc.), which is an analytical quantitative sandwich enzyme immunoassay technique that uses the interaction the mouse monoclonal antibodies with DNA and histone to detect internucleosomal fragmented DNA. The test was performed according to the manufacturer’s instructions.

## Statistical analysis

Results were presented as the means ± standard deviations (SD), the means were compared, and the fold change (effect size) was measured after treatment. Paired observations were compared by paired t-test, hypothesis testing was done using 2-tailed distribution, and 95% confidence intervals were provided for the tests statistics. A p < 0.05 was considered statistically significant for all tests.

## Acknowledgements

This work was supported by Japan Society for the Promotion of Science (JSPS) grant (grant no. 20592328). Special thanks to Dr. Tomohiro Otani and Dr. Yoshimi Takai for helpful discussions.

